# Analytical and Clinical Comparison of Three Nucleic Acid Amplification Tests for SARS-CoV-2 Detection

**DOI:** 10.1101/2020.05.14.097311

**Authors:** Elizabeth Smith, Wei Zhen, Ryhana Manji, Deborah Schron, Scott Duong, Gregory J. Berry

**Affiliations:** Infectious Disease Diagnostics, Northwell Health Laboratories, Lake Success, NY; Department of Pathology and Laboratory Medicine, The Donald and Barbara Zucker School of Medicine at Hofstra/Northwell

**Keywords:** SARS-CoV-2, COVID-19, EUA, molecular diagnostics, TMA, PCR, quantified virus

## Abstract

Severe Acute Respiratory Syndrome Coronavirus 2 (SARS-CoV-2) was first identified in December 2019 and has quickly become a worldwide pandemic. In response, many diagnostic manufacturers have developed molecular assays for SARS-CoV-2 under the Food and Drug Administration (FDA) Emergency Use Authorization (EUA) pathway. This study compared three of these assays: the Hologic Panther Fusion SARS-CoV-2 assay (Fusion), the Hologic Aptima SARS-CoV-2 assay (Aptima) and the BioFire Diagnostics COVID-19 test (BioFire), to determine analytical and clinical performance, as well as workflow. All three assays showed a similar limit of detection (LOD) using inactivated virus, with 100% detection ranging from 500-1,000 genome equivalents/ml, whereas use of a quantified RNA transcript standard showed the same trend, but had values ranging from 62.5 to 125 copies/ml, confirming variability in absolute quantification of reference standards. The clinical correlation found that the Fusion and BioFire assays had a positive percent agreement (PPA) of 98.7%, followed by the Aptima assay at 94.7% when compared to the consensus result. All three assays exhibited 100% negative percent agreement (NPA). Analysis of discordant results revealed that all four samples missed by the Aptima assay had Ct values >37 on the Fusion assay.

In conclusion, while all three assays showed similar relative LODs, we showed differences in absolute LODs depending on which standard was employed. In addition, the Fusion and BioFire assays showed better clinical performance, while the Aptima assay showed a modest decrease in overall PPA. These findings should be kept in mind when making platform testing decisions.

## Introduction

The novel coronavirus was first detected in the United States in Washington State on January 20, 2020 (*1*), and by March 13, 2020, the World Health Organization (WHO) characterized the severe acute respiratory syndrome coronavirus 2 (SARS-CoV-2) outbreak as a worldwide pandemic (*2*). Since then SARS-CoV-2 has rapidly spread across the country. At the time of this report, the number of cases in the U.S. has almost reached 1.4 million, and the number of deaths has exceeded 84,000 according to the Johns Hopkins University database (*3*). Certain regions and cities have much higher prevalence than others, especially the New York metropolitan area that our laboratories serve. New York City alone accounts for approximately 14% of the cases in the United States (*3*).

SARS-CoV-2 is the etiologic agent of coronavirus disease 2019 (COVID-19), for which symptoms can vary from mild (e.g., fever, fatigue, dry cough) to severe illness (e.g., dyspnea, persistent pain or pressure in the chest, hypoxia, confusion), and can ultimately lead to death, particularly for those ≥65 years of age and those with certain underlying medical conditions such as heart disease, lung disease, or diabetes (*4*). Even more challenging, some are infected with the virus but do not display any signs or symptoms of illness, which likely has contributed to the high rate of transmission (R0 = 2.0-2.5) among humans (*5*).

Accurate and sensitive viral detection methods are key to quickly diagnose infections and mitigate transmission. Nucleic acid amplification tests (NAATs) are highly sensitive and specific methods for the detection of SARS-CoV-2 RNA in respiratory specimens. The majority of the SARS-CoV-2 NAAT tests available today are based on real time reverse transcription polymerase chain reaction (RT-PCR) methods, including the BioFire Diagnostics COVID-19 test (BioFire) and the Hologic Panther Fusion SARS-CoV-2 assay (Fusion) evaluated in the present study. Clinical comparative data have been obtained for the Fusion assay (*6, 7, 8, 9*), but to our knowledge there have been no comparative studies of the BioFire assay. Recently, Hologic has developed a second NAAT, the Aptima SARS-CoV-2 assay (Aptima), which has been submitted to the Food and Drug Administration (FDA) for Emergency Use Authorization (EUA); this NAAT is based on target capture and transcription-mediated amplification (TMA) technologies and is run on the Panther instrument. Like the BioFire assay, comparative evaluations have yet to be performed for this new molecular assay.

An increase in testing capability is critically needed to manage the current testing demands, and to monitor and control the spread of the virus going forward. While the FDA has worked quickly to review and authorize more molecular diagnostic platforms to make more options available to meet demand, real world clinical performance and comparative data are still lacking. These data are urgently needed to better understand the benefits and limitations of each test, and how best to incorporate them into local and national testing strategies.

Here we present an analytical and clinical evaluation of these three sample-to-answer NAATs for the qualitative detection of SARS-CoV-2 RNA in symptomatic patients: Fusion, Panther, and BioFire.

## Materials and Methods

### Specimen collection and storage

Nasopharyngeal (NP) swabs were obtained from patients with clinical signs/symptoms of COVID-19. Each collection used a sterile swab made from Dacron, rayon or nylon which was placed into sterile 3 ml universal transport medium (UTM-various manufacturers) after collection. Samples were then transported at room temperature and were stored at 2-8°C for up to 72 hours and tested as soon as possible after collection. For the retrospective sample set, after routine patient testing, aliquots of samples were taken and stored at −80°C until testing of the three comparator NAATs could occur.

### Study design

Samples were selected from specimens received for routine SARS-CoV-2 testing between April and May of 2020. The selection included symptomatic patients of all genders and ages. A total of 150 nasopharyngeal (NP) samples (75 negative and 75 positive) were used for this study: 101 retrospective specimens (51 negative, 50 positive) and 49 prospective, fresh specimens (24 negative, 25 positive). Specimens were selected to represent our laboratory’s positivity rate at the time this study was designed (50 – 60% at beginning of April 2020) and also included positive specimens spanning the range of positivity, including those with low viral loads (characterized by high cycle threshold (Ct) values obtained by results from initial clinical testing). The 101 retrospective specimens were initially tested per routine patient care and then immediately aliquoted and frozen at −80°C, remaining frozen until this study was performed. Retrospective sample aliquots were thawed and immediately tested on the Fusion, Aptima, and BioFire systems. Testing for prospective samples was performed within 48 hours of collection. Prospective samples were run on the Fusion, Aptima and BioFire using the original UTM sample and aliquoted directly into Hologic Lysis tubes and into BioFire Sample Injection Vial. This work was conducted as a quality improvement activity in order to complete assay validation for the Aptima and BioFire assays.

### Hologic Panther Fusion^®^ SARS-CoV-2 assay (Fusion)

The Fusion assay (Hologic, Inc., San Diego, CA) was performed on the Panther Fusion instrument according to the manufacturer’s instructions for use. This assay targets two unique regions of the ORF1ab section of the SARS-CoV-2 viral genome. A 500 μL aliquot of the primary NP swab specimen is transferred into a Specimen Lysis Tube containing 710 μL lysis buffer and is then loaded onto the Panther Fusion instrument. From this tube, the instrument removes 360 μl for extraction. Each specimen is processed with an internal control (IC), which is added via the working Panther Fusion Capture Reagent-S. Nucleic acid is purified using capture oligonucleotides and a magnetic field, eluted in 50 μl, and 5 μl of the eluted nucleic acid is added to a Panther Fusion reaction tube. The Fusion assay amplifies and detects two conserved regions of the ORF1ab section of the SARS-CoV-2 viral genome. Both amplicons are detected by probes using the same fluorescent reporter, with amplification of either or both regions contributing to a single fluorescent signal and single Ct value. Reporting of a positive specimen requires only one of the two targets to be detected (ORF1ab region 1 or ORF1ab region 2).

### Hologic Aptima SARS-CoV-2 assay (Aptima)

The Aptima assay (Hologic, Inc., San Diego, CA) is a NAAT that uses target capture and transcription-mediated amplification (TMA) technologies for the isolation and amplification of SARS-CoV-2 RNA. This assay targets two unique regions of the ORF1ab section of the SARS-CoV-2 viral genome and is performed on the Panther instrument. All testing was performed according to the manufacturer’s instructions and is briefly described. A 500 μL aliquot of the primary NP swab specimen is transferred into a Specimen Lysis Tube containing 710 μL lysis buffer and this tube is then loaded on to the Panther instrument. From the Specimen Lysis Tube, 360 μl is taken for each reaction. Each specimen is processed with an IC, which is added via the working Target Capture Reagent. Nucleic acid is purified using capture oligonucleotides and a magnetic field, and the purified nucleic acid used as template for the TMA reaction. After amplification, chemiluminescent probes hybridize to amplicons and emit light measured by a luminometer in relative light units (RLUs). The IC signal and SARS-CoV-2-specific signal are differentiated by kinetic profiles of the labeled probes (rapid vs slow). Assay results are determined by a cut-off based on the total RLU and the kinetic curve type.

### BioFire COVID-19 Test (BioFire)

The BioFire COVID-19 Test (Salt Lake City, UT) was performed on the BioFire FilmArray 12 Bay Torch System as per the manufacturer’s instructions. This assay targets two unique regions of the ORF1ab section and one unique region of the ORF8 section of the SARS-CoV-2 viral genome. The test pouch (1 per sample to be tested) is prepared by injecting Hydration Solution into the pouch hydration port. Approximately 300 μl of sample is added with a provided transfer pipette to the Sample Injection Vial, followed by the addition of a single-use Sample Buffer Tube to the Sample Injection Vial and inverted to mix. The sample mix is injected into the pouch sample port. Within the pouch, the sample is lysed by agitation (bead beating), and all nucleic acid is extracted and purified using magnetic bead technology. Nested multiplex PCR is performed and endpoint melting curve data is used to detect and generate a result for each target assay on the BioFire COVID-19 Test.

### Analytical Sensitivity

Limit of detection (LOD) was performed using quantified inactivated (gamma-irradiated) SARS-CoV-2 material from Isolate USA-WA1/2020 (NR-52287, BEI Resources, Manassas, VA) and SARS-CoV-2 synthetic RNA quantified control containing five gene targets (E, N, ORF1ab, RdRP and S Genes of SARS-CoV-2) from Exact Diagnostics (SKU COV019, Fort Worth, TX). The BEI material was provided at 4.1 x 10^9 genome equivalents (GE)/ml (2.8 x 10^5 TCID50/ml pre-inactivation of virus) concentration, from which the following serial dilutions were prepared (in GE/ml): 1,000, 500, 250, 125, 62.5, 31.3. Dilutions were prepared using Ambion^®^ RNA Storage Solution (Catalog No. AM7001, ThermoFisher Scientific) to limit the potential of degradation of the RNA, and aliquoted (with replicates ranging from 5-10, as shown in **Table 1**) for testing across the different platforms. The same process was followed for the Exact Diagnostics control, which had starting concentration of 200,000 copies/ml and was diluted to make the same concentration of serial dilutions as those performed for the BEI material. LOD was defined as the lowest dilution at which all replicates were positive (100% detection rate).

**Table 1.** Summary of Limit of Detection results.

|  |  | Positivity Rate (%) <sup>a</sup> |  |  |  |  |  | Final LOD |
| --- | --- | --- | --- | --- | --- | --- | --- | --- |
|  |  | dilution (GE/ml) |  |  |  |  |  |  |
|  |  | 1,000 | 500 | 250 | 125 | 62.5 | 31.3 |  |
| BEI Resources<br>Quantified<br>inactivated virus | Fusion<br>SARS-CoV-2 | **5/5**<br>**(100%)** | 9/10<br>(90%) | 9/10<br>(90%) | 7/10<br>(70%) | 5/10<br>(50%) | 0/10<br>(0%) | 1,000 GE/ml |
|  | Aptima<br>SARS-CoV-2 | 5/5<br>(100%) | **10/10**<br>**(100%)** | 5/10<br>(50%) | 5/10<br>(50%) | 1/10<br>(10%) | 0/10<br>(0%) | 500 GE/ml |
|  | BioFire<sup>b</sup><br>COVID-19 | 5/5<br>(100%) | **10/10**<br>**(100%)** | 9/10<br>(90%) | 7/10<br>(70%) | 5/10<br>(50%) | 0/10<br>(0%) | 500 GE/ml |
|  |  | dilution (copies/ml) |  |  |  |  |  |  |
|  |  | 1,000 | 500 | 250 | 125 | 62.5 | 31.3 |  |
| Exact Diagnostics<br>Quantified<br>synthetic RNA | Fusion<br>SARS-CoV-2 | 5/5<br>(100%) | 10/10<br>(100%) | 10/10<br>(100%) | 10/10<br>(100%) | **10/10**<br>**(100%)** | 8/10<br>(80%) | 62.5 copies/ml |
|  | Aptima<br>SARS-CoV-2 | 5/5<br>(100%) | 10/10<br>(100%) | 10/10<br>(100%) | 10/10<br>(100%) | **10/10**<br>**(100%)** | 8/10<br>(80%) | 62.5 copies/ml |
|  | BioFire<sup>c</sup><br>COVID-19 | 5/5<br>(100%) | 10/10<br>(100%) | 10/10<br>(100%) | **10/10**<br>**(100%)** | 8/10<br>(80%) | 4/5<br>(80%) | 125 copies/ml |

GE, genomic equivalents; LOD, limit of detection
a. The LOD by positivity rate for each NAAT is highlighted in bold.
b. Numbers of replicates at each dilution that gave initial equivocal results and required repeat testing: 0 (1,000 GE/ml), 1 (500 GE/ml), 1 (250 GE/ml), 5 (125 GE/ml), 2 (62.5 GE/ml), 0 (31.3 GE/ml).
c. Numbers of replicates at each dilution that gave initial equivocal results and required repeat testing: 0 (1,000 copies/ml, 500 copies /ml and 250 copies /ml), 2 (125 copies /ml), 6 (62.5 copies /ml), 4 (31.3 copies/ml).

### Workflow Evaluation

Workflow was evaluated using a calibrated timer to measure the time needed for each step being evaluated, including hands-on time (HoT), assay run time and total turnaround time (TAT). HoT, assay run time, and TAT were calculated using the throughput of samples per run.

### Statistical methods

The consensus result was based on the majority results of the 3 NAATs (Fusion, Aptima, BioFire) and was defined as follows: consensus positive = positive result in ≥2 of 3 NAATs; consensus negative = negative result in ≥2 of 3 NAATs. The final result of each NAAT was based on each manufacturer’s results interpretation algorithm. Percent positive agreement (PPA), percent negative agreement (NPA), positivity rate, Kappa, and two-sided (upper/lower) 95% confidence interval (CI) were calculated using Microsoft^®^ Office Excel 365 MSO software (Microsoft, Redmond, WA). As a measure of overall agreement, Cohen’s kappa values (κ) were calculated, with values categorized as follows: >0.90 = almost-perfect, 0.90 to 0.80 = strong, 0.79 to 0.60 = moderate, 0.59 to 0.40 = weak, 0.39 to 0.21 = minimal, 0.20 to 0 = none (10, 11).

## Results

### Analytical Sensitivity

Both quantified inactivated SARS-CoV-2 virus (BEI Resources) and Exact Diagnostics SARS-CoV-2 synthetic RNA quantified control were used to prepare serial dilution panels (1,000 to 31.3 [GE or copies]/ml, in 2-fold dilutions) to determine the LOD of each assay. The LOD was defined as the lowest dilution in which all replicates were detected (100% positivity rate), using the results interpretation algorithm per the instructions for use for each assay. Using these criteria, the LOD using inactivated SARS-CoV-2 virus was 1,000 GE/ml for the Fusion assay, 500 GE/ml for the Aptima assay, and 500 GE/ml for the BioFire test (**Table 1**). In addition, the LOD as determined using the Exact Diagnostics synthetic RNA transcript was 62.5 copies/ml for both the Fusion and Aptima and 125 copies/ml for the BioFire.

### Clinical performance

After testing of the 150 clinical specimens on all 3 platforms, consensus results were determined for each sample (consensus positive = positive result in ≥2 of 3 NAATs; consensus negative = negative result in ≥2 of 3 NAATs) and results from each NAAT were compared to the consensus result. The Fusion and BioFire tests exhibited the highest positive percent agreement (PPA) of 98.7%, while the Aptima assay had a PPA of 94.7% (**Table 2**). NPAs were 100% for all three NAATs, with no false positive results for any platform. Cohen’s kappa values were 0.987, 0.947, and 0.987 for the Fusion, Aptima and BioFire tests, respectively, all of which indicate an “almost perfect” level of agreement with the consensus result.

**Table 2.** Clinical performance comparison of three SARS-CoV-2 NAATs in NP swab specimens (n = 150).

| Molecular Assay | Consensus Result |  | (± 95% CI)<sup>ab</sup> |  |  |
| --- | --- | --- | --- | --- | --- |
|  | Positive | Negative | Kappa (κ) | PPA | NPA |
| Fusion SARS-CoV-2<sup>c</sup> |  |  |  |  |  |
| Positive | 74 | 0 | 0.987 | 98.7% | 100% |
| Negative | 1 | 74 | (0.96 – 1.00) | (0.928 – 0.998) | (0.951 – 1) |
| Aptima SARS-CoV-2 |  |  |  |  |  |
| Positive | 71 | 0 | 0.947 | 94.7% | 100% |
| Negative | 4 | 75 | (0.895 – 0.998) | (0.871 – 0.979) | (0.951 – 1) |
| BioFire COVID-19 |  |  |  |  |  |
| Positive | 74 | 0 | 0.987 | 98.7% | 100% |
| Negative | 1 | 75 | (0.961 – 1.00) | (0.928 – 0.998) | (0.951 – 1) |

a. ±, upper/lower 95%
b. CI, confidence interval
c. One sample gave an invalid result and was excluded from the Fusion agreement analysis.

Initial equivocal or invalid results occurred for 3 samples out of the sample set: 2 equivocal results for the BioFire test and 1 invalid result for the Fusion assay. The initial BioFire results for the 2 samples were one of three targets detected (one of the ORF1ab targets); repeat testing gave the same result for both samples, for an overall interpretation of “detected” as per the manufacturer. The initial Fusion invalid result repeated as invalid and this sample was removed from the overall agreement analysis for the Fusion assay; as this sample was negative by both Aptima and BioFire giving a consensus negative result, this sample was kept in the overall agreement analyses for Aptima and BioFire.

There were no discordant results among the prospective, fresh sample set; all of the discordant samples occurred among the retrospective sample set. Of the three NAATs evaluated, the Aptima assay had the most discordant results with the consensus (n=4), followed by the Fusion and BioFire tests each with one discordant result. Discordant sample results are shown in **Table 3**. The four discordant samples for Aptima had Fusion Ct values ≥37.3, indicating lower viral titers in these samples. The discordant BioFire sample had a Fusion Ct value of 35.7. For the 5 Fusion/Aptima discordant samples, the BioFire COVID-19 Test detected only 2 of 3 targets in each sample (GSD-3, GSD-6, GSD-23) or detected 1 of 3 targets in each sample twice for an overall result of detected (GSD-4, GSD-48) (**Table 3**).

**Table 3.** Details of discordant samples.

| SAMPLE ID | Individual SARS-CoV-2 NAAT results<sup>a</sup> |  |  |  |
| --- | --- | --- | --- | --- |
|  | Consensus Result | Fusion SARS-CoV-2 (Ct value) | Aptima SARS-CoV-2 | BioFire COVID-19 |
| **GSD-3** | positive | positive (38.1) | **NEGATIVE** | detected (2d/2e) |
| **GSD-4** | positive | positive (38.1) | **NEGATIVE** | equivocal (2a) twice, overall detected |
| **GSD-6** | positive | positive (40.5) | **NEGATIVE** | detected (2a/2e) |
| **GSD-7** | positive | positive (35.7) | positive | **NOT DETECTED** |
| **GSD-23** | positive | **NEGATIVE** (N/A) | positive | detected (2a/2e) |
| **GSD-48** | positive | positive (37.3) | **NEGATIVE** | equivocal (2a) twice, overall detected |

<sup>a</sup> Discordant sample results are highlighted in bold

### Workflow

Workflow parameters along with basic assay characteristics are presented in **Table 4**. Maximum sample throughput in 8 hour and 24 hour time periods were considerably higher on the Fusion and Aptima platforms (Panther Fusion: 335/8 hours and 1150/24 hours; Panther Aptima: 275/8 hour and 1020/24 hours), while throughput for BioFire was 72/ 8 hour and 216/ 24 hours. Hands on time (HoT) per specimen for the both Fusion and Aptima assays are 1 minute, where the BioFire HoT is 3 minutes per specimen. Sample preparation for the Fusion and Aptima assays consist of aliquoting 500 μL of sample into a lysis tube. Sample preparation for the BioFire consists of five steps including pouch hydration, sample and buffer mixing, addition of sample/buffer to the pouch, and loading onto the instrument. These additional step for BioFire account for the increased HoT per specimen as seen in **Table 4**.

**Table 4.** Basic assay characteristics and workflow parameters of three EUA SARS-CoV-2 NAATs

|  | **Hologic Panther Fusion**<br>**SARS-CoV-2 Assay** | **Hologic Aptima**<br>**SARS-CoV-2 Assay** | **BioFire**<br>**COVID-19 Test** |
| --- | --- | --- | --- |
| Detection platform/System | Panther Fusion | Panther or Panther Fusion | BioFire FilmArray Torch<br>System—12 bay tower |
| Sample type<sup>a</sup> | NP, OPS, LRT, NS | NP, OPS,<br>NS, nasal wash/aspirate | NP |
| Sample volume required (μL) | 500 μl (250 μl for LRT) | 500 μl | 300 μl |
| Target region of SARS-CoV-2 | two regions of ORF1ab | two regions of ORF1ab | two regions of ORF1ab<br>and ORF8 |
| Analytical sensitivity per claim | 0.01 TCID<sub>50</sub>/ml | 0.01 TCID<sub>50</sub>/ml | 0.022 TCID<sub>50</sub>/ml<br>(330 genomic copies /ml) |
| Observed analytical sensitivity -<br>**Inactivated Virus** | 1000 genomic copies/ ml | 500 genomic copies/ ml | 500 genomic copies/ ml |
| Observed analytical sensitivity -<br>**RNA Transcript** | 62.5 copies/ ml | 62.5 copies/ ml | 125 copies/ ml |
| High throughput processing | Yes | Yes | No |
| Throughput- Samples per run | 120<br>with continuous loading | 120<br>with continuous loading | 12 |
| Hands on Time/ sample | 1 minute | 1 minute | 3 minutes |
| Hands on Time/Run | 2 hours / 120 samples | 2 hours / 120 samples | 36 minutes / 12 samples |
| Assay Processing Time/ Run | 4 hours 35 minutes /<br>120 samples | 5 hours 30 minutes /<br>120 samples | 45 minutes / 12 samples |
| Time to first result | 2 hours 25 minutes | 3 hours 30 minutes | 45 min |
| Overall Turn Around Time/<br>Run | 6 hours 35 minutes /<br>120 samples | 7 hours 30 minutes /<br>120 samples | 1 hour 21 minutes /<br>12 samples |
| Maximum Sample throughput<br>in 8/24 hrs | 335 / 1150 | 275 / 1020 | 72 / 216 |

a. NP, nasopharyngeal swab; NS, nasal swab; OPS, oropharyngeal swab; LRT, lower respiratory tract.

## Discussion

The present study compared the analytical sensitivity (LOD), clinical performance, and workflow of three SARS-CoV-2 NAATs in 150 NP swab specimens. Our two independent LOD analyses revealed that while all 3 assays had an LOD that was within 1 dilution factor of each other within a given control material, absolute LODs with quantified inactivated virus (500-1,000 GE/ml) were several-fold higher than the value obtained when using quantified synthetic RNA (62.5-125 copies/ml). This difference in absolute LOD values reflects the inherent difficulty in comparing standards that have been prepared and quantitated differently and have very high stock concentrations requiring significant dilutions for LOD panel testing (i.e., any quantification error and/or pipetting error of the stock will be magnified in a dilution series). Of interest, the LOD calculations for BioFire using the synthetic RNA standard only showed amplification in two of three gene targets, since ORF1ab is included in the Exact diagnostics control, but ORF 8 is not included. Considering this discrepancy, the decreased LOD relative to the Fusion and Aptima when using the quantified synthetic RNA is likely due to the lack of ORF 8 target material. Our clinical correlation showed that the Fusion and Biofire had similar PPA (98.7%), while the Aptima showed a slight decrease at 94.7% PPA due to 4 false negative results among samples in the frozen retrospective set. Cohen’s kappa values showed almost perfect agreement (range: 0.947 – 0.987) between each assay and the consensus result. NPAs were 100% for all NAATs, suggesting each of these assays has high specificity.

All discordant results were false negatives as compared to the consensus result, with Fusion and BioFire each demonstrating one missed positive and Aptima demonstrating four missed positives. A closer analysis of discordant results showed that the sample missed by the Fusion (GSD-23) only had two gene targets detected by BioFire (2a/2e) and the sample missed by the BioFire (GSD-7) had a Ct value of 35.7 on the Fusion. All four samples missed by the Aptima (GSD-3, 4, 6, and 48) had Ct values ranging from 37.3-40.5, suggesting that the Aptima assay is slightly less sensitive than the Fusion and Biofire assays. Of note, two of the specimens missed by the Aptima (GSD-4 and 48) were also each equivocal twice by BioFire and resulted as positive as per the EUA IFU, suggesting that these were also weak positives on the BioFire assay.

Workflow comparisons between the Fusion and the Aptima were quite similar, both being optimal in high-volume testing situations (> 1,000 tests/24 hours), whereas the BioFire assay had an advantage of fast sample to answer time, allowing for faster detection and diagnosis that is useful in urgent situations.

Other comparator studies of the Fusion assay have recently been reported, including: a study comparing Fusion to the modified CDC assay, GenMark ePlex, DiaSorin Simplexa (6); a study comparing Cepheid, ID NOW, and GenMark using Fusion as the reference standard (7); a study comparing Fusion, Cepheid, DiaSorin, cobas, and a laboratory-developed test (LDT) (8); and a study comparing Fusion to an in-house LDT targeting the envelope gene (9). In general, these reports show that the Fusion SARS-CoV-2 assay is comparable, if not superior, to other molecular platforms available for SARS-CoV-2 testing. In particular, these evaluations showed that Fusion, DiaSorin and Cepheid tended to have better analytical sensitivity and fewer false negatives in clinical testing than other assays. The low rate of false negatives is a critical performance aspect, as missed positive results allow for further spread of the disease and potential patient mismanagement. These comparison studies, while not clinical trials as would be typical of new FDA-cleared IVD assays, have provided a window into the relative real world clinical performance characteristics of these assays, which has been a significant knowledge gap since these SARS-CoV-2 EUA molecular assays first became available. Among these knowledge gaps are relative performance data for the Aptima and BioFire tests. To our knowledge, this is the first report comparing both the Aptima and BioFire assay to any other NAAT for SARS-CoV-2 detection.

It is worth noting that the LOD for the Fusion SARS-CoV-2 assay as determined in this study was 1,000 GE/ml when using inactivated virus, while the LOD determined in a previous study by our laboratory was 83 ±36 copies/ml (6) and an additional LOD study performed as part of this study once again using the same synthetic quantified RNA standard also previously published showed a comparable LOD of 62.5 copies/ml. This is an important additional new set of data that exhibits the difficulty of comparing absolute LOD values when using different standards, specifically inactivated virus vs. synthetic RNA standards.

This study does have limitations, being a single site study with a limited number of NP swab specimens (n = 150) included in the clinical evaluation. However, this specimen set was selected to be representative of our patient population (including samples from patients of all ages and both genders) and positivity rate (50-60% in the beginning of April 2020), and included samples with a range of viral loads (low, moderate, high). Some initial equivocal or invalid results (n=3) by BioFire and Fusion required retesting, but only 1 remained unresolved on retest and was removed from the agreement analysis for the Fusion assay.

In conclusion, our data show that the Fusion, Aptima and BioFire SARS-CoV-2 NAATs exhibit similar LODs, even when tested using two different quantified standards. In addition, the Fusion and BioFire assays have comparable clinical performance for detection of SARS-CoV-2 in NP swabs, with a PPA of 98.7%, while the Aptima assay showed a slight reduction at 94.7% PPA. All 3 assays demonstrated 100% NPA, suggesting high specificity. These performance characteristics, as well as testing volume and workflow requirements, should be considered when making testing platform decisions.

## Acknowledgements and Disclosures

Gregory Berry has previously given education seminars for Hologic, Inc. and BioFire Diagnostics and has received Honorariums.

